# Histone H3K4me1 and H3K27ac play roles in nucleosome depletion and eRNA transcription, respectively, at enhancers

**DOI:** 10.1101/2021.01.05.425373

**Authors:** Yujin Kang, Yea Woon Kim, Jin Kang, AeRi Kim

## Abstract

Histone H3K4me1 and H3K27ac are enhancer specific modifications and are required for enhancers to activate transcription of target genes. However the reciprocal effects of these histone modifications on each other and their roles in enhancers are not clear. Here to comparatively analyze the role of these modifications, we inhibited H3K4me1 and H3K27ac by deleting SET domain of histone methyltransferases MLL3 and MLL4 and HAT domain of histone acetyltransferase p300, respectively, in erythroid K562 cells. The loss of H3K4me1 reduced H3K27ac at the β-globin enhancer LCR HSs, but H3K27ac reduction did not affect H3K4me1. This unequal relationship between two modifications was revealed in putative enhancers by genome-wide analysis using ChIP-seq. Histone H3 depletion at putative enhancers was weakened by the loss of H3K4me1 but not by the loss of H3K27ac. Chromatin remodeling complexes were recruited into the β-globin LCR HSs in a H3K4me1-dependent manner. In contrast, H3K27ac was required for enhancer RNA (eRNA) transcription, and H3K4me1 was not enough for it. Forced H3K27ac induced eRNA transcription without affecting H3K4me1 at the β-globin LCR HSs. These results indicate that H3K4me1 and H3K27ac affect each other in different ways and play more direct roles in nucleosome depletion and eRNA transcription, respectively, at enhancers.

## INTRODUCTION

Enhancers are cis-regulatory elements that highly induce the transcription of target genes (Plank and Dean 2014; Smith and Shilatifard 2014). They have unique histone modifications, H3K4me1 and H3K27ac, in the chromatin environment. H3K4me1 and H3K27ac are catalyzed by histone methyltransferases MLL3 and MLL4 (Herz et al. 2012; Hu et al. 2013) and histone acetyltransferases CBP and p300 (Jin et al. 2011; Hilton et al. 2015), respectively, in mammalian cells. Depletion or inactivation of these histone modifying enzymes disturbs cell-type-specific gene expression in addition to the severe decrease of H3K4me1 or H3K27ac at enhancers (Kim and Kim 2013; Lee et al. 2013; Wang et al. 2016; Ebrahimi et al. 2019). However, the reciprocal effects of these histone modifications on each other at enhancers are not clear. Most studies have focused on one of the histone modifications by targeting histone modifying enzymes, and controversial results have been obtained from different experimental approaches such as knockout of the modifying enzyme genes, mutation in the catalytic domains and chemical inhibition of the activity domains (Lee et al. 2013; Dorighi et al. 2017; Rickels et al. 2017; Raisner et al. 2018).

Enhancers have a special chromatin structure that is hypersensitive to attack by endonucleases such as DNase I and MNase, which allows access of transcription factors (TFs) to their binding motifs (Guertin and Lis 2013; Inukai et al. 2017). Histones are depleted at the hypersensitive enhancers as revealed by ChIP assay (Fang et al. 2009). Studies using MNase-treatment have shown the lack of nucleosome structure at DNase I hypersensitive sites (HSs) (Kim et al. 2007; Grossman et al. 2018). When DNase I sensitivity is increased at enhancers by transcriptional activation, it accompanies more severe depletion of nucleosomes (Johnson et al. 2018). Enhancers defined by H3K4me1 are hypersensitive to DNase I attack (Zentner et al. 2011). These findings suggest a correlation of enhancer specific histone modifications with nucleosome or histone depletion.

Long non-coding RNAs are transcribed from enhancers by RNA polymerase II (pol II), which are referred to as enhancer RNAs (eRNAs) (Kim et al. 2010). eRNA transcription is activated by transcription factors such as p53 and nuclear hormone receptor, and by Mediator (Lai et al. 2013; Li et al. 2013; Melo et al. 2013). Transcribed eRNAs have been reported to contribute to many transcriptional activation steps including the recruitment of RNA pol II to promoters (Mousavi et al. 2013; Maruyama et al. 2014), chromatin interaction between enhancers and promoters (Li et al. 2013; Hsieh et al. 2014) and the elongation transition of paused RNA pol II (Schaukowitch et al. 2014). eRNAs-producing enhancers have specific histone modifications including H3K27ac (Zhu et al. 2013). Depletion of MLL3/4 or treatment with a CBP/p300 bromodomain inhibitor reduce eRNA levels, suggesting the correlation of H3K4me1 and/or H3K27ac with eRNAs transcription (Rahnamoun et al. 2018; Raisner et al. 2018).

Here, to analyze how enhancer-specific histone modifications H3K4me1 and H3K27ac affect each other and what is the distinct roles of these modifications in enhancer activity, we inhibited H3K4me1 and H3K27ac by deleting the SET domain of MLL3 and MLL4 (ΔMLL3/4) and HAT domain of p300 (Δp300), respectively, using the CRISPR/Cas9 system in erythroid K562 cells. H3K4me1 and H3K27ac were comparatively and reciprocally examined at the β-globin enhancer locus control region (LCR) HSs in ΔMLL3/4 and Δp300 K562 cells (Fig. 1A). In order to expand results from the β-globin locus into the whole genome, we carried out ChIP-sequencing. Histone H3 occupancy was analyzed at the regulatory regions including enhancers in ΔMLL3/4 cells and Δp300 cells. Recruitment of chromatin remodeling complexes was examined at cell specific enhancers including the β-globin LCR HSs. The roles of H3K4me1 and H3K27ac in eRNA transcription were explored by analyzing total RNA-seq data. These studies indicate that H3K4me1 and H3K27ac affect each other in different ways and play distinct roles in activating enhancers.

**Figure 1.**
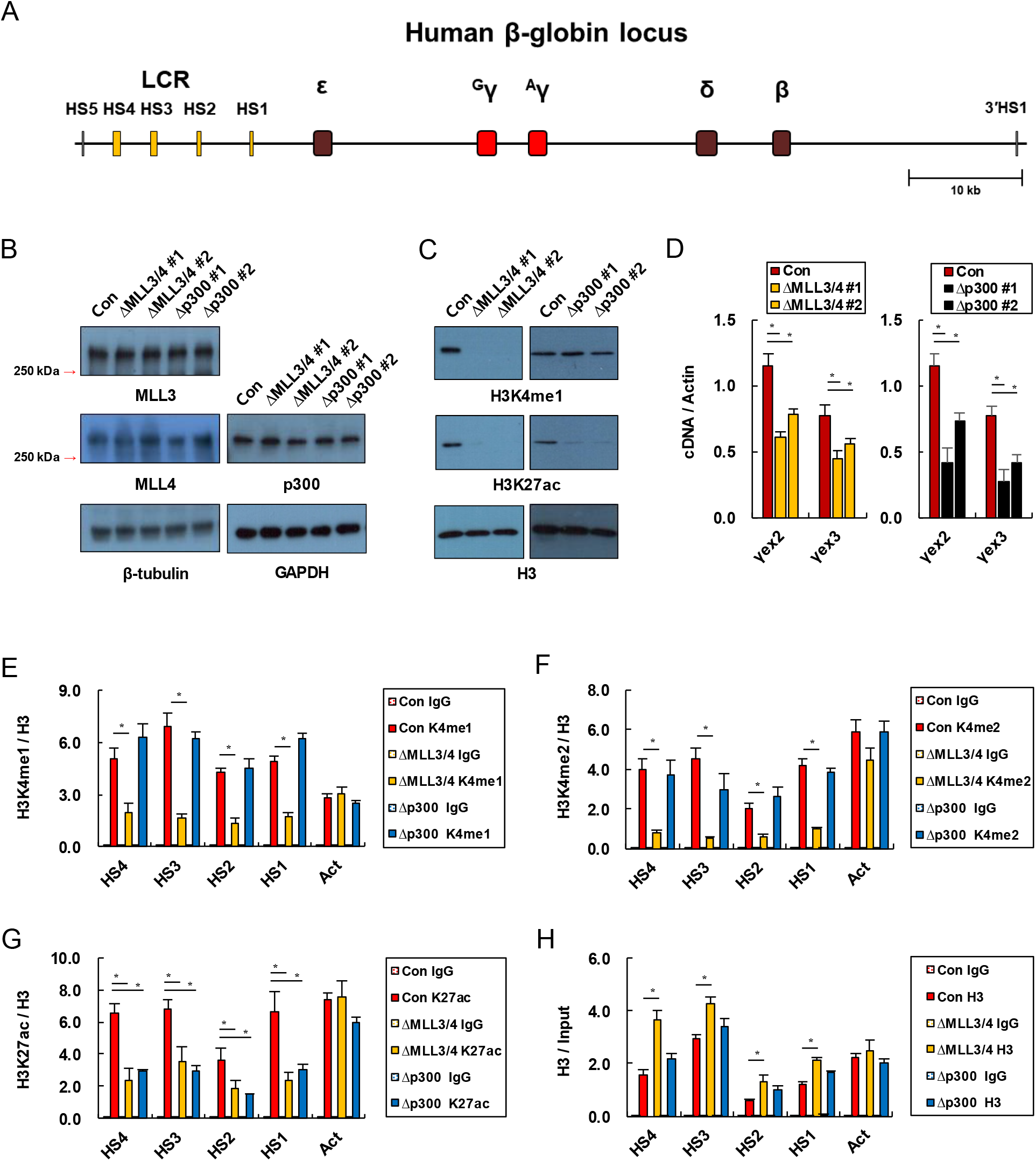
H3K4me1 and H3K27ac at the β-globin enhancer LCR HSs in ΔMLL3/4 cells and Δp300 K562 cells. (A) The diagram represents the human β-globin locus. The β-globin LCR HSs and β-like globin genes are indicated by yellow and red/brown rectangles, respectively. Black vertical bars represent insulators. Protein levels of MLL3, MLL4 and p300 (B) and histone H3K4me1 and H3K27ac (C) were analyzed by western blot in control cells (Con) and two clones of ΔMLL3/4 cells and Δp300 cells. The β-tubulin, GAPDH and H3 ware used as loading control. (D) Transcription of the human γ-globin genes were measured by quantitative RT-PCR. The amounts of cDNA for the γ-globin genes were compared with a genomic DNA standard and then normalized to the Actin cDNA. The results of 4 to 7 independent experiments were graphed with ± SEM. Enrichments of histone H3K4me1 (E), H3K4me2 (F) and H3K27ac (G) were measured using ChIP assay and quantitative PCR at the β-globin LCR HSs. DNA immunoprecipitated by antibodies to H3K4me1, H3K4me2 and H3K27ac were quantitatively compared with DNA reacted by H3 antibodies. (H) DNA immunoprecipitated by H3 antibodies was compared with input DNA. The Actin was used as positive control. Normal rabbit IgG (IgG) served as negative experimental control. Results are presented as the means ± SEM of 3 to 5 independent experiments. The p-values were calculated using the two tailed Student’s t-test (*P < 0.05).

## RESULTS

### Loss of H3K4me1 decreases H3K27ac at the β-globin enhancer LCR HSs, but loss of H3K27ac does not affect H3K4me1

To study reciprocal effects of enhancer specific histone modifications H3K4me1 and H3K27ac on each other, we deleted DNA sequences encoding catalytic domains of MLL3 (KMT2C) and MLL4 (KMT2D) or p300 using CRISPR/Cas9 technique in human erythroid cell line K562 (Supplemental Fig. S1). Deletions of SET domains in MLL3/4 (ΔMLL3/4) and deletion of HAT domain in p300 (Δp300) did not affect protein stability of MLL3, MLL4 and p300 when it was analyzed by western blotting with antibodies against N-terminal regions (Fig. 1B). Amounts of histone H3K4me1 and H3K27ac were decreased by ΔMLL3/4 and Δp300, respectively, at the total protein levels (Fig. 1C). While Δp300 did not affect amount of H3K4me1, H3K27ac was reduced in two ΔMLL3/4 clonal cells in spite of unaffected p300 expression. The loss of catalytic domains reduced transcription of the erythroid specific γ-globin genes in K562 cells (Fig. 1D). To analyze enhancer histone organization, we carried out ChIP assay and examined the β-globin enhancer LCR using qPCR. Histone H3K4me1 and H3K4me2 were decreased by ΔMLL3/4 at the LCR HSs, but not by Δp300 (Fig. 1E, F). H3K27ac was decreased by ΔMLL3/4 in addition to Δp300 (Fig. 1G). These results imply that H3K4me1 affects H3K27ac in enhancers, but not the reverse. In addition, ChIP analysis for histone H3 indicated that H3K4me1 contributes to histone depletion at the β-globin LCR HSs (Fig. 1H).

### Genome wide analysis shows H3K4me1 is required for H3K27ac in putative enhancers but H3K27ac is not for H3K4me1

In order to expand observations from the β-globin LCR HSs into the whole genome, we carried out ChIP-seq analysis for H3K4me1 and H3K27ac. Our ChIP-seq data in control cells were consistent with public ENCODE ChIP-seq data in K562 cells (Supplemental Fig. S2). 108,210 regions were sorted by H3K4me1 enrichment in control cells (Fig. 2A). H3K4me1 was almost completely eliminated by ΔMLL3/4 in 90% (n = 96,928) of the regions, where H3K27ac was also severely reduced. In the other hand, Δp300 decreased H3K27ac in 79% (n = 89,199) of this modification peaks (n = 112,261) (Fig. 2B), even though the decreases were not severe as the decreases of H3K4me1 by ΔMLL3/4. H3K4me1 was maintained in regions where H3K27ac is decreased. These results indicate that ΔMLL3/4 and Δp300 inhibit H3K4me1 and H3K27ac, respectively, genome-wide. H3K4me1 appears to affect H3K27ac, but it is not likely to be the reverse case.

**Figure 2.**
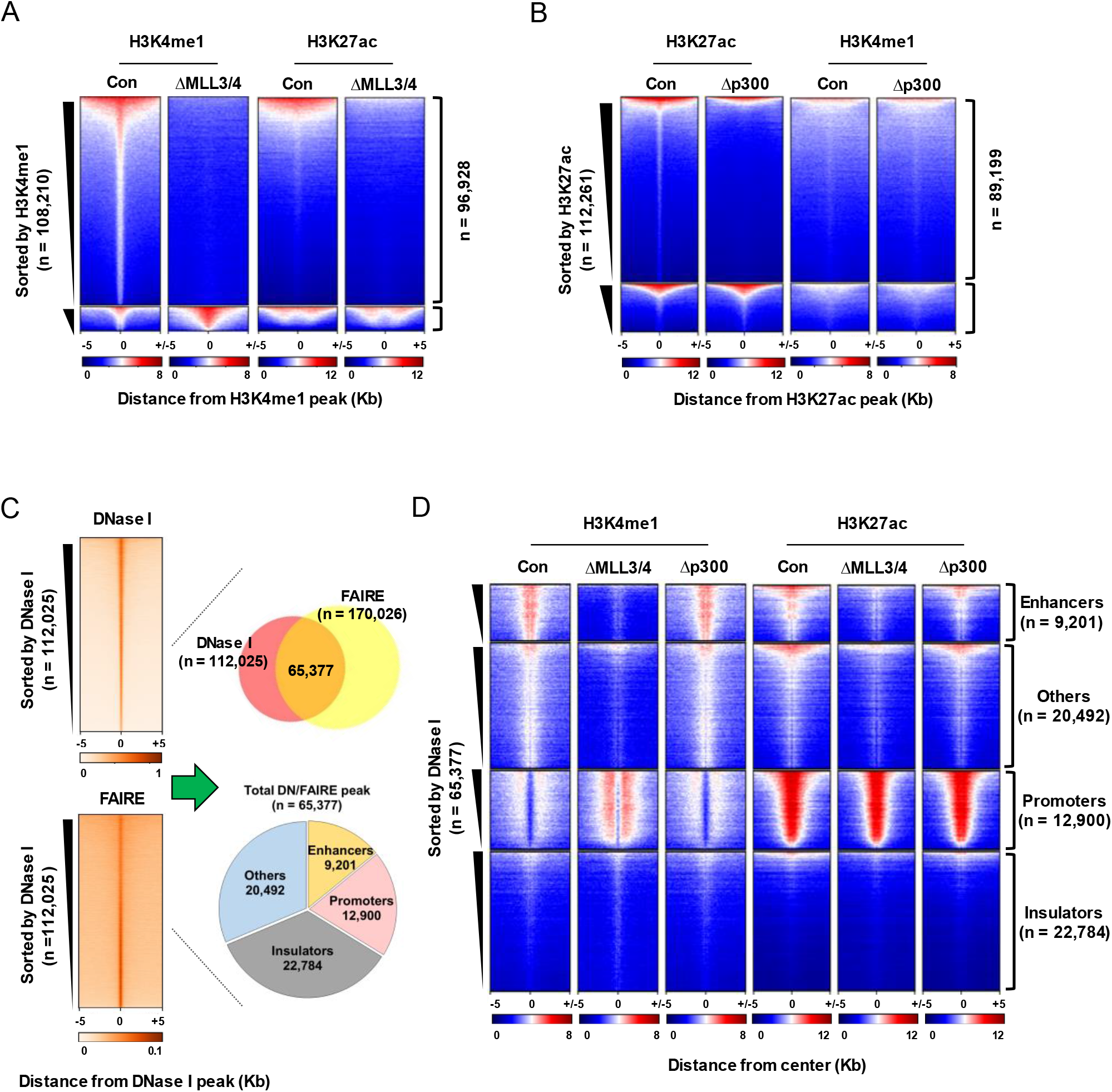
Genome wide analysis of H3K4me1 and H3K27ac in ΔMLL3/4 cells and Δp300 cells. (A) Heatmaps show signal enrichment of H3K4me1 and H3K27ac at H3K4me1 peaks in control cells (Con) and ΔMLL3/4 cells. H3K4me1 regions (n = 108,210) were divided into two groups; H3K4me1-reduced regions in ΔMLL3/4 cells (0 < Con CPM −ΔMLL3/4 CPM, n = 96,928) and others (0 ≥ Con CPM – ΔMLL3/4 CPM, n = 11,282). (B) Signal enrichment of H3K27ac and H3K4me1 was showed at H3K27ac peaks in Con and Δp300 cells by heatmaps. H3K27ac regions (n = 112,261) were divided into two groups; H3K27ac-reduced regions in Δp300 cells (0 < Con CPM – Δp300 CPM, n = 89,199) and others (0 ≥ Con CPM – Δp300 CPM, n = 23,062). Five kilobase pairs around the center of H3K4me1 or H3K27ac are displayed. Color scales indicate the relative signal intensity on heatmaps. (C) DNase I and FAIRE enrichment were presented by heatmaps at DNase I peak (± 5 Kb from the center) in K562 cells. Venn diagram depicts overlapping regions in DNase-seq and FAIRE-seq peaks (n = 65,377, right top panel). The overlapping regions were classified into four groups; enhancers, promoters, insulators and others. (D) H3K4me1 and H3K27ac ChIP signal (± 5 Kb from the center) were presented by heatmaps at the four groups.

To focus on the relationship between H3K4me1 and H3K27ac at enhancers, we have screened HSs by overlapping DNase-seq data and FAIRE-seq data (Fig. 2C). The HSs (n = 65,377) were classified into putative enhancers by H3K4me1 in extragenic regions, promoters by TSS ± 1 Kb, insulators by CTCF occupancy and the ‘others’ for the remaining regions. H3K4me1 was most notable in putative enhancers, where H3K4me1 was not dependent on H3K27ac, but H3K27ac was dependent on H3K4me1 (Fig. 2D). Unexpectedly, H3K4me1 was increased in promoter by ΔMLL3/4 and this might relate with ratio to H3K4me3. Modification patterns in the ‘others’ were relatively similar with the patterns in putative enhancers. This might be because the ‘others’ include enhancers located in genes/introns. Thus, this genome-wide analysis shows that H3K4me1 is required for H3K27ac in enhancers, but it is not likely to be in the opposite direction. It is consistent with the results obtained from the β-globin enhancer LCR HSs.

### Histone depletion at enhancers is dependent on H3K4me1 but not on H3K27ac

Our results from the β-globin LCR HSs suggested that H3K4me1 might be required for histone depletion in enhancers (Fig. 1G). To study this possibility, we analyzed histone H3 occupancy genome-widely in ΔMLL3/4 cells and Δp300 cells and then compared it at the HSs screened by DNase-seq data and FAIRE-seq data. Histone H3 was depleted at the HSs as shown by heatmap (Fig. 3A). The depletion was reduced by the loss of H3K4me1 at putative enhancers and ‘others’ that may include enhancers (Fig. 3A, B). Reduction of H3K27ac did not significantly affect H3 depletion at the HSs including putative enhancers. To study the correlation between H3K4me1 and histone depletion at enhancers, the putative enhancers were sorted by H3K4me1 level and divided into three groups according to the levels (Fig. 3C). Comparison between the groups showed that H3K4me1 level parallels histone H3 depletion. It also correlates with chromatin accessibility by ATAC-seq data and with nucleosome depletion by MNase-seq data (Fig. 3D), supporting H3K4me1-dependent histone depletion at enhancers.

**Figure 3.**
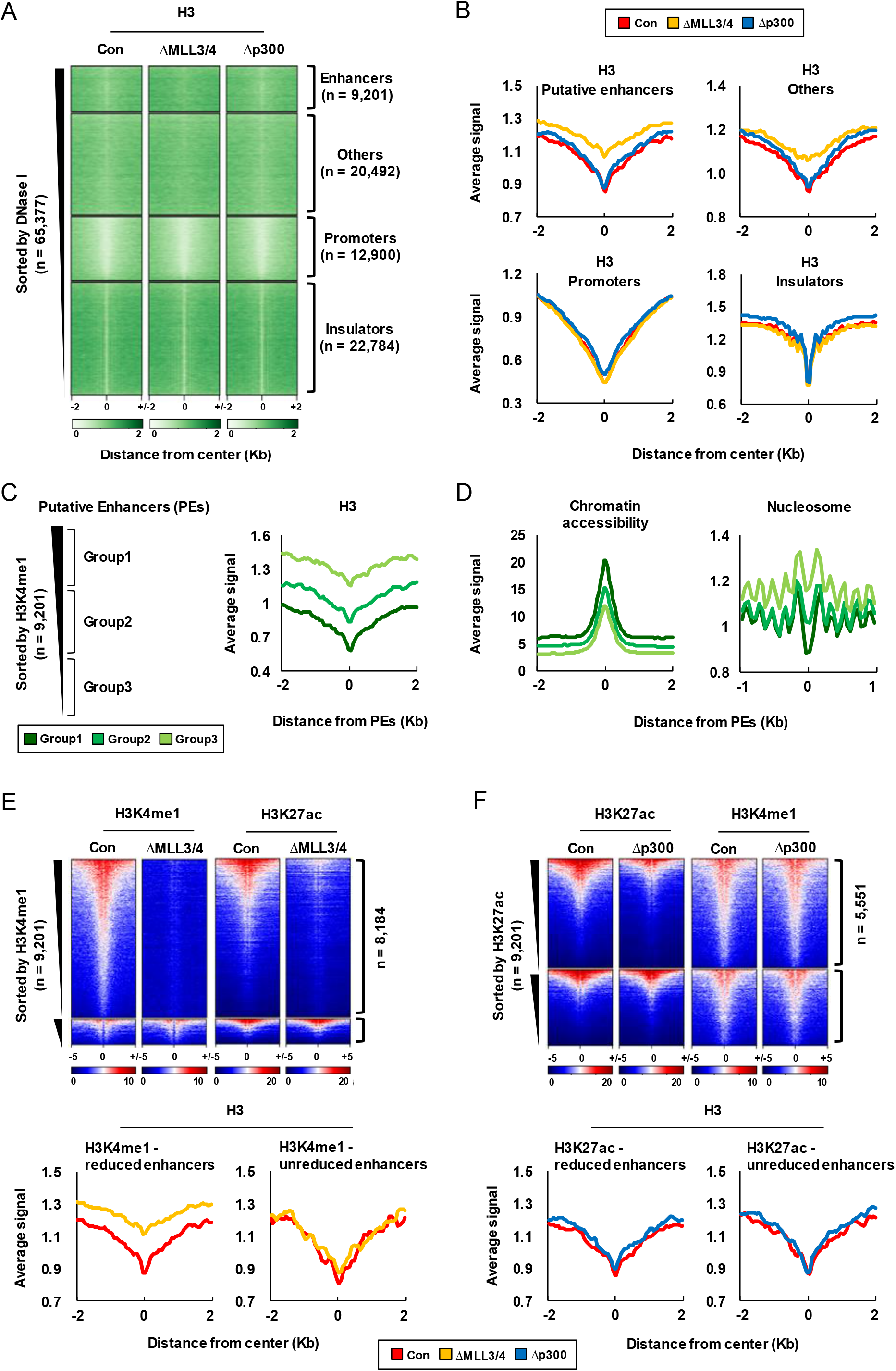
Histone depletion at enhancers in ΔMLL3/4 cells and Δp300 cells. Histone H3 ChIP signal (± 2 Kb from the center) was presented by heatmaps for four groups classified in Fig. 2C (A) and by average profiles (B). (C) Putative enhancers (PEs, n = 9,201) were sorted by H3K4me1 signal in Con cells and divided into three groups according to the signal (left). Average profiles of H3 signal (± 2 Kb from the PEs center) were drawn for the groups (right). (D) Average profiles of chromatin accessibility signal (± 2 Kb from the PEs center, left) and nucleosome depletion signal (± 1 Kb from the PEs center, right) were drawn using public ATAC-seq and MNase-seq data. (E) Heatmaps present signal enrichment of H3K4me1 and H3K27ac (± 5 Kb from the PEs center) in two PEs groups; H3K4me1-reduced in ΔMLL3/4 cells (0 < Con CPM – ΔMLL3/4 CPM, n = 8,184) and H3K4me1-unreduced (0 ≥ Con CPM – ΔMLL3/4 CPM, n = 1,017). H3 signal (± 2 Kb from the center) in two PEs groups was presented by average profiles (lower). (F) Heatmaps present signal enrichment of H3K27ac and H3K4me1 (± 5 Kb from the PEs center) in two PEs groups; H3K27ac-reduced in Δp300 cells (0 < Con CPM – Δp300 CPM, n = 5,551) and H3K27ac-unreduced (0 ≥ Con CPM – Δp300 CPM, n = 3,650). H3 signal (± 2 Kb from the center) in two PEs groups was presented by average profiles (lower).

To more clarify relationship of H3K4me1 and H3K27ac with histone depletion at enhancers, we divided putative enhancers into two groups depending on the changes of H3K4me1 by ΔMLL3/4 (Fig. 3E). Histone H3 depletion was more weakened in H3K4me1-reduced enhancers (n = 8,184), even it was not in the rest enhancers where H3K4me1 was not reduced. In contrast, histone depletion was maintained in enhancers where H3K27ac was apparently reduced by Δp300 (n = 5,551) (Fig. 3F). H3K4me1 was also maintained in the H3K27ac-reduced enhancers. Similar histone depletion patterns were observed in ‘Others’ regions by reduction of H3K4me1 or H3K27ac (Supplemental Fig. S3). Taken together, these results indicate that H3K4me1 is required for histone depletion at enhancers. H3K27ac does not appear to directly contribute to the histone depletion.

### Chromatin remodeling complexes are recruited into enhancer in a H3K4me1-dependent manner

Because histones/nucleosomes can be depleted by the ATP-dependent chromatin remodeling process, we aligned public ChIP-seq data for chromatin remodeling complexes to putative enhancers. BRG1 (SMARCA4), INI1 (SMARCB1) and BAF170 (SMARCC2), subunits of SWI/SNF (BAF) chromatin remodeling complex, and SNF2h (SMARCA5), a subunit of ACF chromatin remodeling complex, were robustly detected at the enhancers (Fig. 4A) and their levels positively correlated with histone H3K4me1 level (Supplemental Fig. S4). Views of genome browser showed these subunits bind to the β-globin LCR HSs (Fig. 4B). In order to explore the role of H3K4me1 in recruiting chromatin remodeling complexes, we performed ChIP assay using antibodies for BRG1, SNF2h and ACF1 (BAZ1A, a subunit of ACF complex). These subunits were less detected at the β-globin LCR HSs in ΔMLL3/4 cells compared to control cells (Fig. 4C-E). Other erythroid specific enhancers (Kang et al. 2015), where the subunits bind to (Supplemental Fig. S5), showed similar binding patterns with the LCR HSs. The results suggest a positive role of H3K4me1 in recruitment of chromatin remodeling complexes. However, no significant changes were observed in Δp300 cells. Genes coding the remodeling complex subunits were similarly expressed in control cells, ΔMLL3/4 cells and Δp300 cells (Fig. 4F). Thus, these results indicate that H3K4me1 plays a role in recruiting chromatin remodeling complexes into enhancers, but H3K27ac is not necessary for it.

**Figure 4.**
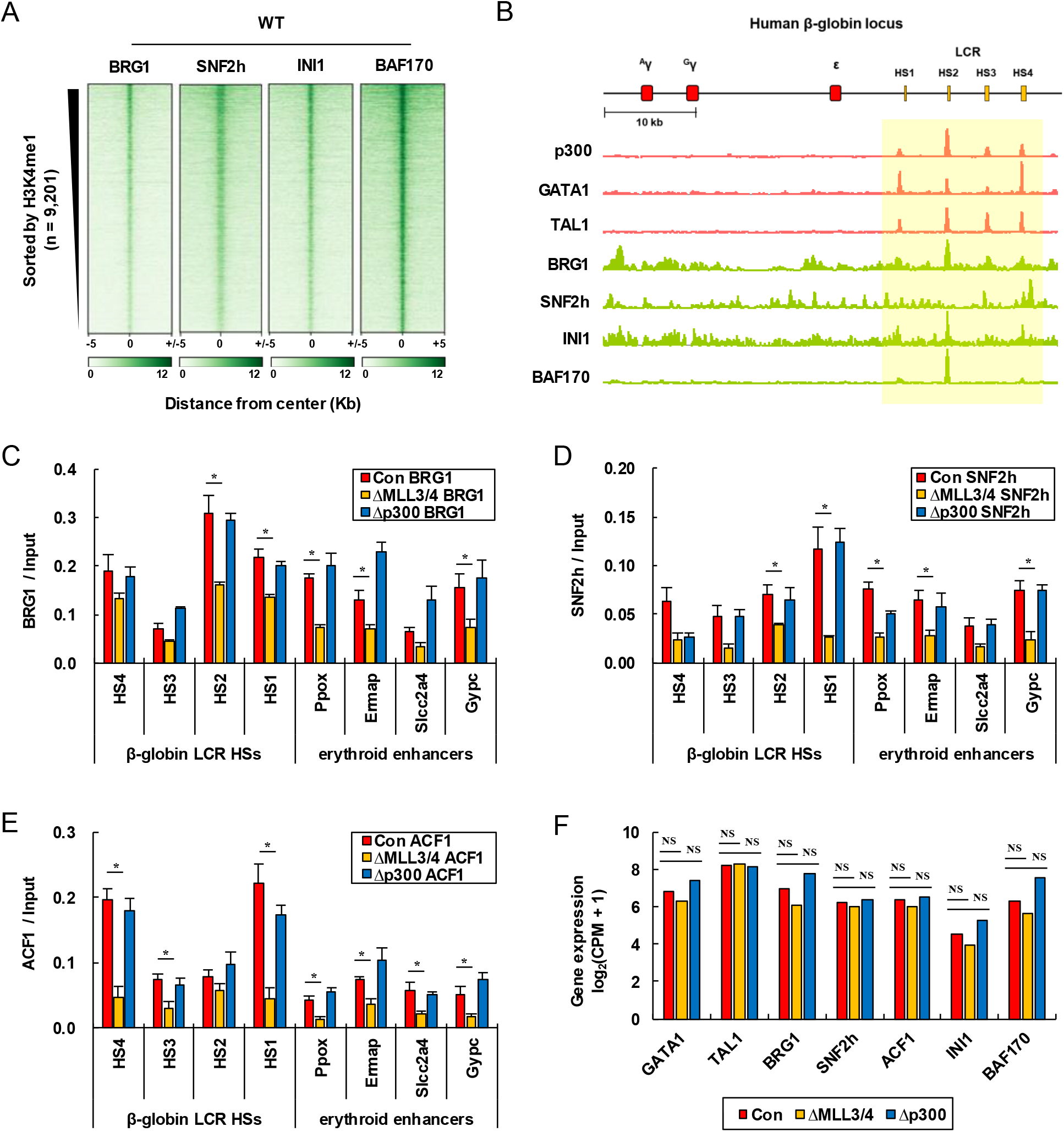
Occupancy of subunits of chromatin remodeling complexes at enhancers in ΔMLL3/4 cells and Δp300 cells. (A) BRG1, SNF2h, INI1 and BAF170 signal (± 5 Kb from the center) were presented by heatmaps at PEs in K562 cells (WT). (B) IGB genome browser tracks show distribution of p300, GATA1, TAL1, BRG1, SNF2h, INI1 and BAF170 at the β-globin locus. Enhancers are highlighted with yellow shadow. The occupancy of BRG1 (C), SNF2h (D) and ACF1 (E) was analyzed by ChIP assay at the β-globin enhancers and other erythroid specific enhancers in Con cells, ΔMLL3/4 cells and Δp300 cells. The amount of immunoprecipitated DNA was quantitatively compared with input DNA. The results of 4 independent experiments were graphed with ± SEM. The p-values were calculated using the two tailed Student’s t-test (*P < 0.05). (F) Expression levels of GATA1, TAL1, BRG1, SNF2h, ACF1, INI1 and BAF170 were determined using total RNA-seq data. The y axis indicates log2-transformed (CPM + 1).

### eRNA transcription requires histone H3K27ac in addition to H3K4me1

To study the roles of enhancer specific histone modifications in eRNA transcription, we divided putative enhancers into active and poised enhancers according to the levels of H3K27ac as shown by heatmap (Fig. 5A) and measured transcription in the enhancer regions. eRNAs were more highly transcribed in active enhancers than poised enhancers. eRNA transcription from active enhancers was reduced in ΔMLL3/4 cells and Δp300 cells (Fig. 5B). Transcription of nearest genes (n = 500) from eRNAs-decreased enhancers was also reduced as revealed by Gene Set Enrichment Analysis (GSEA) (Fig. 5C). These results indicate that H3K27ac is required for eRNA transcription in addition to H3K4me1.

**Figure 5.**
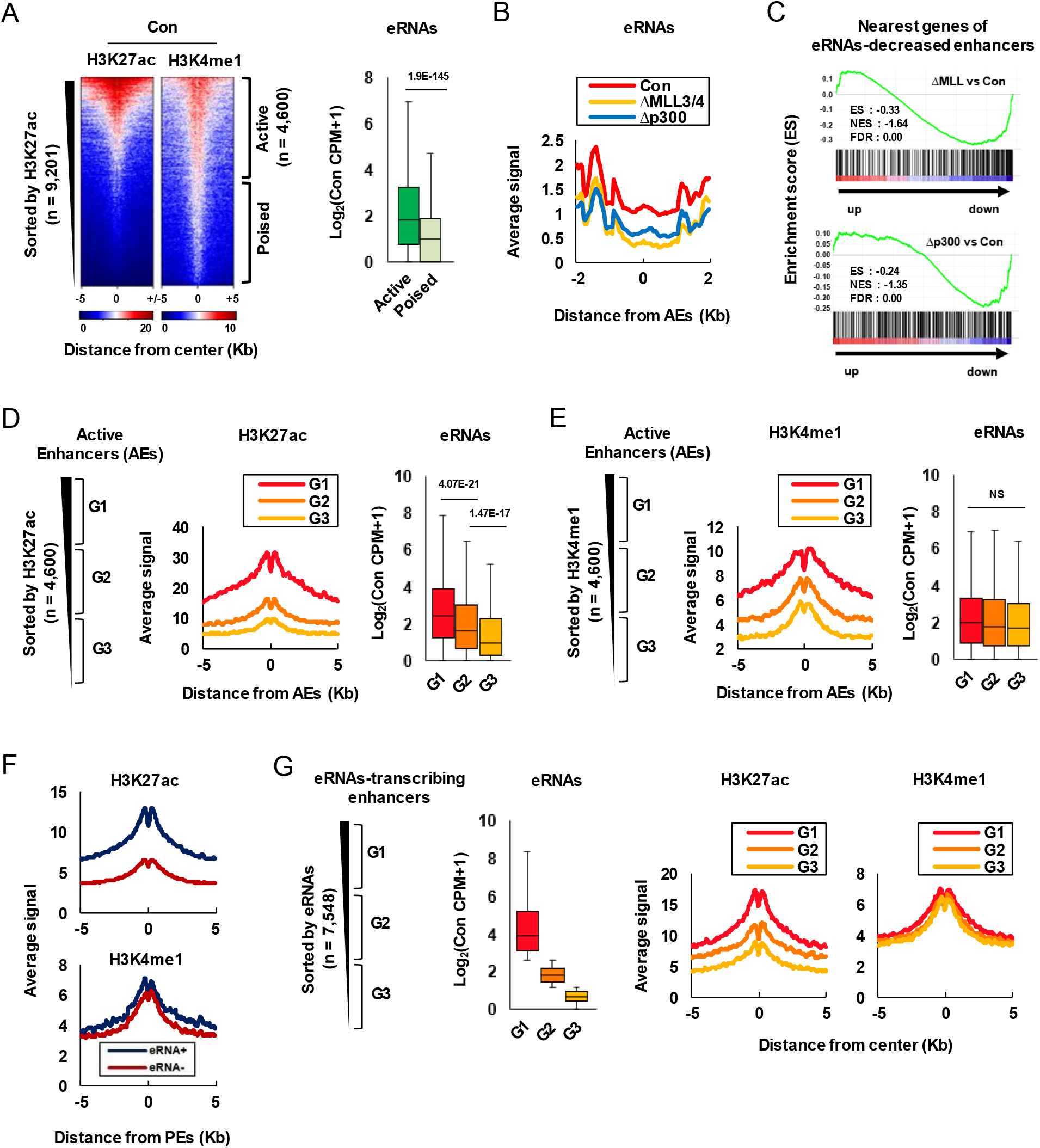
eRNA transcription in ΔMLL3/4 cells and Δp300 cells. (A) PEs were divided into active (n = 4,600) and poised (n = 4,601) according to H3K27ac signal in Con cells. Signal enrichment of H3K27ac and H3K4me1 (± 5 Kb from the center), and eRNA transcription were presented using heatmaps and boxplot, respectively, for both enhancer groups. (B) eRNA expression (± 2 Kb from the center) from active enhancers (AEs) was depicted by average profiles in Con cells, ΔMLL3/4 cells and Δp300 cells. (C) Expression for the nearest genes of eRNA-decreased AEs in ΔMLL3/4 cells (upper) and in Δp300 cells (lower) was analyzed by gene set enrichment analysis (GSEA). ‘Up’ and ‘down’ indicate the relative transcription level of genes in ΔMLL3/4 cells and Δp300 cells compared to Con cells. AEs were divided into three groups according to H3K27ac signal (D) and H3K4me1 signal (E) in Con cells. Each histone modification signal and eRNA transcription of the groups were presented in average profile and boxplot, respectively. (F) PEs (n = 9,201) were divided into two groups according to eRNA transcription; eRNA+ (0 < Con eRNAs CPM, n = 7,548) and eRNA- (0 = Con eRNAs CPM, n = 1,653). Average profiles represent H3K27ac and H3K4me1 signals in two enhancer groups. (G) eRNA-transcribing enhancers (n = 7,548) were divided into three groups according to the eRNA signal in Con cells. Boxplot depicts expression level of eRNAs at the groups. Average signals of H3K27ac and H3K4me1 (± 5 Kb from the center) were profiled on each groups. The y axis of all boxplots indicates log2-transformed (CPM + 1).

To further analyze relationship of H3K27ac and H3K4me1 with eRNA transcription, active enhancers (n = 4,600) were divided into three groups according to H3K27ac levels or H3K4me1 levels in control cells and transcription levels were measured in each groups (Fig. 5D, E). eRNA levels positively correlate with H3K27ac, but there was no significant correlation between eRNA levels and H3K4me1 at active enhancers. When putative enhancers were divided depending on eRNA transcription, H3K27ac was higher in enhancers transcribing eRNAs (n = 7,548) than not transcribing them, but there were no great differences in H3K4me1 between the enhancer groups (Fig. 5F). Analysis according to eRNA levels showed that eRNA transcription correlates with H3K27ac strongly but does so with H3K4me1 at a very weak level (Fig. 5G). Thus this analysis supports that H3K27ac plays a role in eRNA synthesis at enhancers prepared by H3K4me1.

### Increase of H3K27ac by TSA treatment induces eRNA transcription at the β-globin LCR HSs without increase of H3K4me1

To determine whether H3K27ac has a direct role in eRNA transcription, we treated mouse erythroid MEL/ch11 cells with histone deacetylase inhibitors TSA. A human chromosome 11 containing the β-globin locus is present in MEL/ch11 cells. TSA treatment for 6 h and 24 h increased H3K27ac at the total protein level, without notable changes on H3K4me1 (Fig. 6A). It also elevated H3K27ac at the chromatin level as shown by ChIP assay in the β-globin enhancer LCR HSs (Fig. 6B). Surprisingly, eRNA levels were increased at the LCR HSs with 6 h and 24h TSA treatment (Fig. 6C), and it brought about transcription of β-like globin genes following eRNA transcription (Fig. 6D). However, H3K4me1 was largely maintained through the β-globin locus (Fig. 6E). Thus, these results indicate that histone H3K27ac plays a more direct role in eRNA transcription from enhancers rather than H3K4me1.

**Figure 6.**
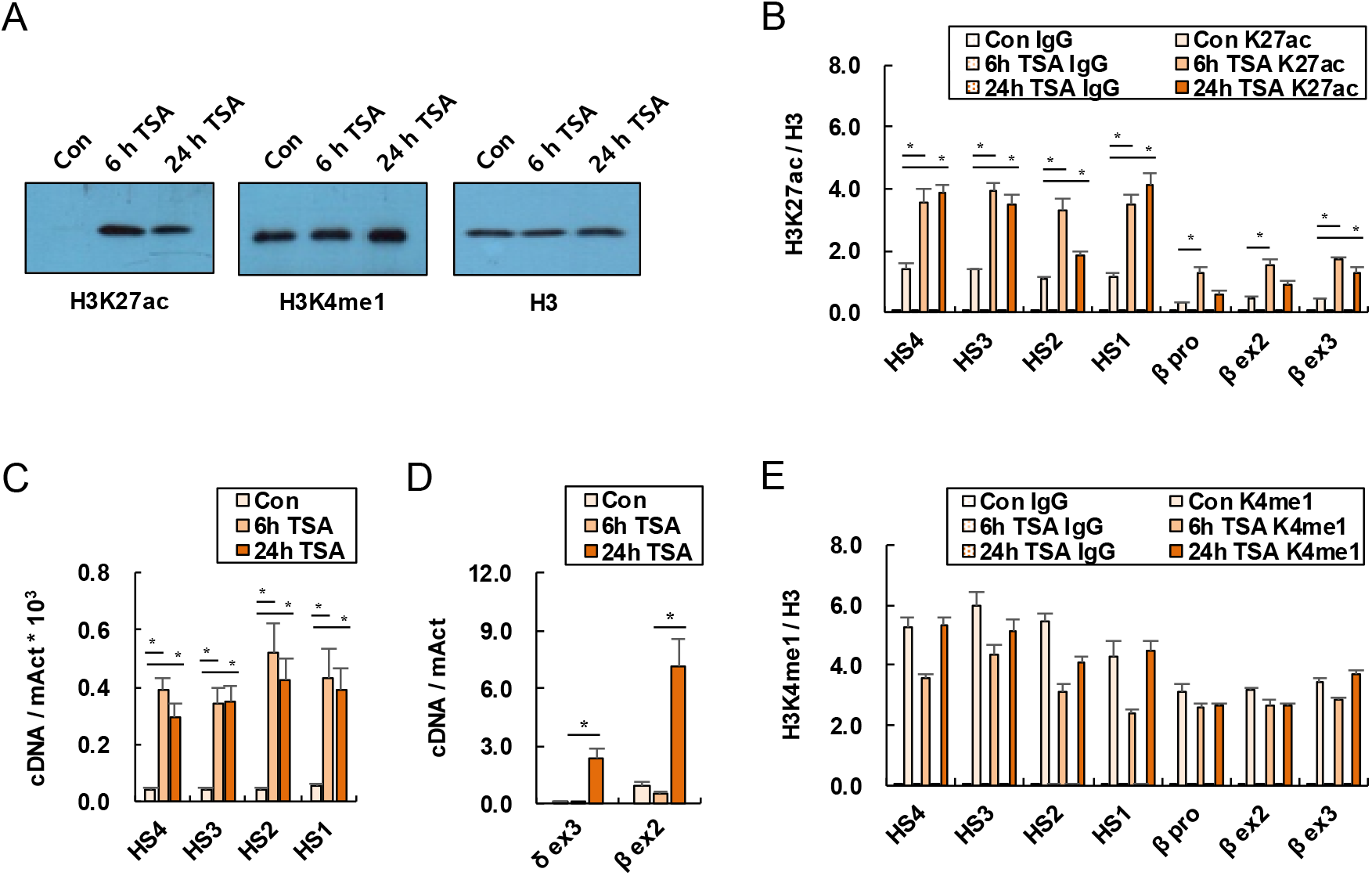
eRNA transcription from the β-globin LCR HSs in TSA-treated MEL/ch11 cells. (A) Protein levels of H3K27ac, H3K4me1 and H3 were detected by Western blot in control and TSA-treated cells. The H3 was used as experimental control. H3K27ac (B) and H3K4me1 (E) at chromatin level were analyzed by ChIP assay in the β-globin LCR HSs and the promoter and exons of the β-globin gene. eRNA from the LCR HSs (C) and mRNA from the β-like globin genes (D) were measured by quantitative RT-PCR. The results of 4 to 5 independent experiments were graphed with ± SEM. The p-values were calculated using the two tailed Student’s t-test (*P < 0.05).

## DISCUSSION

Our comparative and reciprocal studies show that histone H3K4me1 and H3K27ac affect each other at enhancers in a different way; H3K4me1 is required for H3K27ac but this is not true in the opposite direction (Fig. 7). H3K4me1 appears to play a role in histone depletion at enhancers by recruiting chromatin remodeling complexes. H3K27ac is not likely to be necessary for the histone depletion. Instead, eRNA transcription seems to require H3K27ac in addition to H3K4me1. Thus our results suggest that enhancer specific histone modifications H3K4me1 and H3K27ac play distinct roles in activating enhancers.

**Figure 7.**
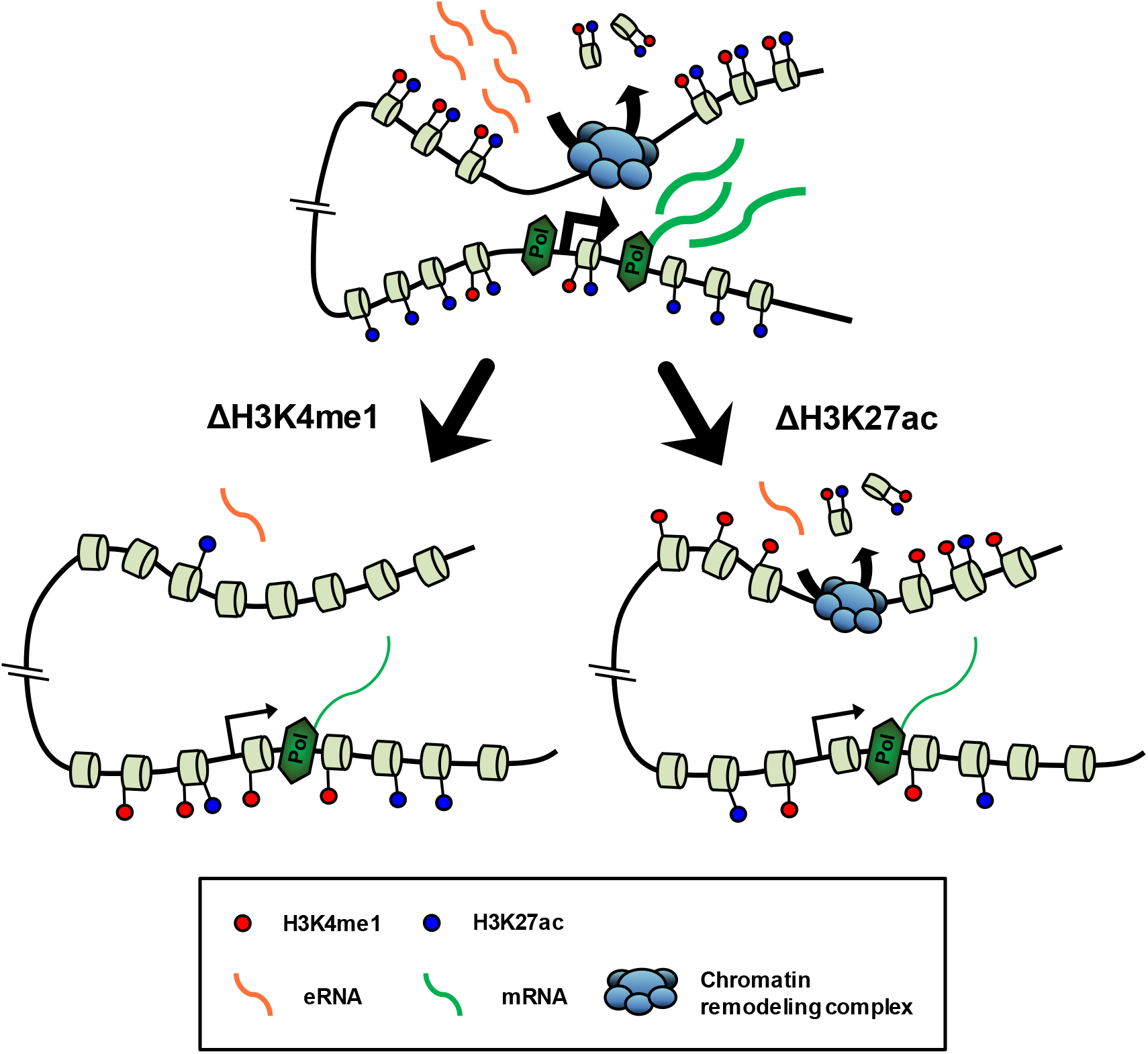
Effects by depletion of H3K4me1 and H3K27ac on enhancers. Results from all Figures were combined and represented in diagrams. Depletion of H3K4me1 at enhancers impairs H3K27ac, histone depletion, recruitment of chromatin remodeling complexes, eRNA transcription and mRN A transcription of target gene. Depletion of H3K27ac at enhancers decreases eRNA transcription and its target gene transcription without affecting H3K4me1, histone depletion and recruitment of chromatin remodeling complexes.

Results from the β-globin locus and genome-wide analysis show that H3K4me1 is required for H3K27ac at enhancers, but H3K27ac is not for H3K4me1. This might be because H3K4me1 precedes H3K27ac at poised enhancers but H3K27ac is deposited at active enhancers. There are reports that MLL3/4, rather than H3K4me1, are important for p300 recruitment and H3K27ac (Wang et al. 2016; Dorighi et al. 2017) and MLL3 and MLL4 are destabilized by deletion of the catalytic domains (Jang et al. 2019). However, our catalytic domain deletion did not affect the stability of MLL3 and MLL4 and eliminated H3K4me1 at enhancers, resulting in reduction of H3K27ac. Similar effects were reported in mouse ES cells where MLL3/4 catalytic domain deletion resulted in reduction of H3K4me1 and H3K27ac at active enhancers without affecting stability and occupancy of MLL3 and MLL4 (Rickels et al. 2017). These findings support the contribution of H3K4me1 to H3K27ac at enhancers. The effects of H3K27ac on H3K4me1 are controversial in many studies using chemical inhibitors to CBP/p300 activity (Wang et al. 2017; Raisner et al. 2018; Zhu et al. 2018; Ebrahimi et al. 2019). However, our study using p300 catalytic domain mutation indicates that H3K27ac is not necessary for H3K4me1 at enhancers, even though depletion of H3K27ac was not severe as the depletion of H3K4me1 by ΔMLL3/4. Similar results were observed in a study using CBP/p300 bromodomain inhibitor, where H3K27ac was reduced at enhancers but H3K4me1 was maintained (Raisner et al. 2018). Taken together, we conclude that the effects of H3K4me1 and H3K27ac on each other at enhancers are not reciprocal.

Histone H3K4me1 appears to play a role in recruiting chromatin remodeling complexes into enhancers, resulting in histone depletion. Our analysis showed that several subunits of the complexes co-localize with enhancers selected by H3K4me1 and are recruited in a H3K4me1-dependent manner. BRG1, a subunit of SWI/SNF (BAF) chromatin remodeling complex, has been reported to regulate DNase I sensitivity and histone H3 occupancy at enhancers of the α-globin locus (Kim et al. 2009). Depletion of BRG1 impaires the extension of nucleosome linker regions by nucleosome shifting at tissue specific enhancers (Hu et al. 2011). Consistent with our finding, binding of BRG1 is decreased by loss of H3K4me1 in the Sox2 enhancer (Local et al. 2018). H3K4me1-containing mononucleosomes are more efficiently bound and remodeled by BAF complex than H3K4me3-containing mononucleosomes in *in vitro* assay (Local et al. 2018). In addition, DPF3, a subunit of BAF complex, has been reported to recognize H3K4me1 using the PHD1-PHD2 domain (Lange et al. 2008; Local et al. 2018). The PHD domain is also present in ACF1 subunit (Eberharter et al. 2004) that interacts with SNF2h in ACF chromatin remodeling complex (He et al. 2006). Thus H3K4me1 may contribute to histone depletion by providing recognition sites for subunits of chromatin remodeling complexes.

When it is compared to H3K4me1, H3K27ac appears to play a more direct role in eRNA synthesis, even though these two modifications are required for it. eRNA transcription is reduced by inhibition of H3K27ac via CBP/p300 bromodomain inhibitor (Raisner et al. 2018) or by loss of H3K4me1 via depletion of MLL3/4, even though it can be due to the loss of MLL3/4 rather than the loss of H3K4me1 (Dorighi et al. 2017; Rahnamoun et al. 2018). However, in our studies, H3K4me1 was maintained at enhancers when H3K27ac is inhibited by Δp300, implying that H3K27ac is more critical for eRNA transcription and H3K4me1 is not sufficient. This proposal is supported by analysis using ChIP-seq data and RNA-seq data that show a positive correlation of eRNA transcription with H3K27ac but no significant correlation with H3K4me1 (Fig. 5). H3K27ac can be recognized by BRD4 (Filippakopou lo s et al. 2012; Roe et al. 2015). BRD4 has been reported to be required for recruiting RNA pol II into enhancers (Nagarajan et al. 2014; Lee et al. 2017). Thus H3K27ac is thought to contribute to eRNA transcription by recruiting RNA pol II into enhancers, where H3K4me1 might be already deposited.

## METHODS

### Cell culture and Trichostatin A treatment

K562 cells were grown in RPMI 1640 medium, and 293FT and MEL/ch11 cells were cultured in DMEM medium as previously described (Kang et al. 2017). MEL/ch11 cells (5 × 10^5^ cells/ml) were treated with 25 ng/ml Trichostatin A (TSA, Sigma) for 6 hours (h) and 24 h in complete DMEM medium.

### Deletion of catalytic domain of MLL3/4 or p300 using CRISPR/Cas9 system

Single guide RNA (sgRNA) sequences were designed for SET domains of MLL3 and MLL4 and HAT domain of p300 with online tools (http://crispr.mit.edu). Two complementary oligos containing gRNA sequences and flanking sequences for cloning were phosphorylated and annealed as previously described (Kim and Kim 2017). Sequences of oligos for sgRNAs are presented in Supplemental Table S1. Annealed oligos for MLL4 SET or p300 HAT were inserted into lentiCRISPRv2 vector (Addgene #52961) (Sanjana et al. 2014) and annealed oligos for MLL3 SET were inserted into pLH-spsgRNA2 vector (Addgene #64114) (Ma et al. 2015). The vectors were cloned in Stbl3 bacteria, prepared using the Plasmid Midi Kit (Qiagen), and transfected into 293FT cells using the Virapower packaging mix (Invitrogen) and Lipofectamine 2000 (Invitrogen). Lentiviruses for MLL4 SET and p300 HAT sgRNA were harvested after 3 days and transduced into K562 cells in the presence of 6 μg/ml polybrene. The transduced cells were selected by 2 μg/ml puromycin and seeded in 96 well plates to obtain clones. Lentivirus for MLL3 SET sgRNA was transduced into MLL4 SET deletion K562 cells and was selected by 500 μg/ml hygromycin. Genomic DNA flanking the MLL3/4 SET domain or p300 HAT domain was amplified by PCR (Supplemental Fig. S1). Primer sequences are listed in Supplemental Table S2.

### Western blot analysis

Proteins were extracted from 2 × 10^6^ cells using RIPA buffer with sonication. Equal amounts of the proteins were electrophoresed in 4-15% SDS-PAGE gradient gel (Bio-rad) and transferred to 0.2 μm NC membrane or 0.45 μm PVDF membrane. After blocking with 5% skim milk in PBST, the membrane was incubated with primary antibodies against H3 (ab1791), H3K4me1 (ab8895), and H3K27ac (ab4729) from Abcam and p300 (sc-8981) and β-tubulin (sc-9104) from Santa Cruz Biotechnology and MLL3 and MLL4 from Jaehoon Kim (KAIST, Korea) at 4 °C overnight and then with secondary antibodies, anti-rabbit HRP (sc-2030) in PBST with 1% skim milk. Rabbit polyclonal anti-MLL3 and anti-MLL4 antibodies were developed against purified histidine-tagged human MLL3 and MLL4 fragments (amino acid residues 1-120 and 4442-4561, respectively), and affinity purified (AbClon). Protein signals were exposed using ECL reagent.

### Reverse transcription-PCR (RT-PCR)

Total RNA was extracted from 2 × 10^6^ cells using QIAzol Lysis Reagent (Qiagen) and cDNA was generated from 0.5 μg of RNA with random hexamers using the GoScript kit (Promega) as previously described (Kang et al. 2018). Quantitative PCR was performed with TaqMan probes in 10 μl of reaction volume using a 7300 Real-time PCR system (Applied Biosystems). The sequences of the probes and primers are listed in Supplemental Table S3.

### Chromatin immunoprecipitation (ChIP)

ChIP was performed as previously described (Cho et al. 2008). Cells (1 × 10^7^) were cross-linked in 1% formaldehyde and digested using MNase. Fragmented chromatin was reacted with antibodies and recovered using protein A or G agarose beads. DNA was purified by phenol extraction and ethanol precipitation and then analyzed by quantitative PCR. The sequences of primers for ChIP assay are presented in Supplemental Table S3, S4. Antibodies used for ChIP experiment were H3 (ab1791), H3K4me1 (ab8895), H3K27ac (ab4729), SNF2h (ab3749) from Abcam, H3K4me2 (07-030) from Millipore, ACF1 (A301-318A) from Bethyl Laboratories and BRG1 (sc-17796) from Santa Cruz Biotechnology. Normal rabbit IgG (sc-2027) from Santa Cruz Biotechnology was used as a negative control.

### RNA library preparation and sequencing

Total RNA was extracted using RNeasy Plus Mini Kit (Qiagen) according to the manufacturer’ s instructions and qualified using Qubit Fluorometer (RNA IQ > 7). After depleting ribosomal RNA (rRNA) using NEBNext rRNA Depletion Kit (New England Biolabs #E6350L), RNA was reversely transcribed using NEBNext Ultra II Directional RNA Library Prep Kit (New England Biolabs #E7765S) as suggested by the manufacturer. This procedure includes fragmentation, priming using random primers, first-strand cDNA synthesis, second-strand cDNA synthesis, cDNA end repair, adaptor ligation, amplification using adaptor primers (8 cycles), and purification using beads. Final libraries were quantified by Qubit dsDNA HS assay (Invitrogen) and 100 bases paired-end reads were sequenced using an Illumina NovaSeq 6000 system.

### RNA-seq analysis

Quality scores of sequenced reads were assessed using quality control tool. Average score was Q = 38 at each base across reads. Input read ends with poor quality values were removed from raw reads by Trimmomatic (quality score 20) (Bolger et al. 2014). Remaining reads were aligned to the human reference genome (hg19) using STAR (Dobin et al. 2013). Exon annotation GTF file was obtained from UCSC database. Aligned reads were counted per gene ID (Entrez) using featureCounts (Liao et al. 2014). Differentially Expressed Genes (DEG) were statistically analyzed using edgeR (Robinson et al. 2010) with q-value under 0.05. CPM values above 1 were considered as meaningful transcription level in genes. Gene set enrichment analysis (GSEA) was performed according to the instructions at GSEA platform (www.gsea-msigdb.org/gsea/index.jsp). Genes near enhancers were sought using GREAT platform (http://great.stanford.edu/public/html) and limited to 500 genes by eRNA reduction.

### ChIP DNA library preparation and sequencing

ChIP DNA (10 ng for H3K4me1, H3K27ac, H3 and input) was processed with NEBNext Ultra II DNA Library Prep Kit (New England Biolabs #E7103S) according to manufacturer’s instructions. ChIP DNA were repaired at the ends, ligated with 1.5 μM NEBNext adaptors, selected in 200 bp size with NEBNext sample purification beads, and amplified with the adaptor primers (7 cycles). After purifying, final libraries were quantified with Qubit dsDNA HS assay (Invitrogen) and 100 bases paired-end reads were sequenced on an Illumina NovaSeq 6000 system.

### ChIP-seq analysis

ChIP-seq raw reads were qualified by removing input reads with poor quality values (quality score 20) and then aligned to the hg19 canonical genome using Bowtie2 (Langmead and Salzberg 2012). Aligned BAM files were filtered by minimum MAPQ quality score 20 and sorted by chromosomal coordinate. Potential PCR duplicates were removed from BAM files. Peak regions of H3K4me1 and H3K27ac were identified using MACS2 (Feng et al. 2012) providing histone H3 data as control. Thresholds of enrichment q-value were 0.05 for narrow peak and 0.05 for broad peak. Matrix files were generated from BAM files and bigwig files using ComputeMatrix, and then used for plotting heatmaps and average profiles with plotHeatmap and plotProfile tools, respectively (Ramírez et al. 2016). Bin size of matrix files was 50 bp. ChIP-seq signals were visualized using Integrated Genome Browser (IGB) (Freese et al. 2016).

### Public NGS data

Public NGS data were obtained from Gene Expression Omnibus (GEO). GEO accession numbers of the data are H3K4me1_1 (GSM788085), H3K4me1_2 (GSM733692), H3K27ac_1 (GSM733656), H3K27ac_2 (GSM646434), p300 (GSM935401), GATA1 (GSM 1003608), TAL1 (GSM935496), BRG1 (GSM935633), SNF2h (GSM2424122), INI1 (GSM935634), BAF170 (GSM3634134), DNase-seq (GSM816655), FAIRE-seq (GSM864361), ATAC-seq (GSM3452726) and MNase-seq (GSM920557) in K562 cells and HeLa RNA-seq (GSM765402) in HeLa-S3 cells.

### Statistical analysis

All qPCR results are presented as the means ± standard error of the mean (SEM) of 3 to 5 independent experiments. The p-values were calculated using the two tailed Student’s t-test (*P < 0.05). Boxplot data is statistically analyzed using two tailed Student’s t-test between two groups of classified enhancers. All boxplots indicate log2-transformed (CPM + 1).

## DATA ACCESS

All raw and processed sequencing data generated in this study have been submitted to the NCBI Gene Expression Omnibus (GEO; https://www.ncbi.nlm.nih.gov/geo/) under accession number GSE147826.

## ACKNOWLEDGEMENTS

We are grateful to Jaehoon Kim for kind gifts of antibodies against MLL3 and MLL4. This research was supported by Basic Science Research Program through the National Research Foundation of Korea (NRF) funded by the Ministry of Science, ICT and Future Planning (NRF-2014R1A2A1A11051702), and by PNU-RENovation (2019-2020).

## AUTHOR’S CONTRIBUTIONS

Y. Kang and A. Kim designed research; Y. Kang performed research; Y. Kang analyzed data; Y. Kang, Y.W. Kim, J. Kang and A. Kim discussed results; Y. Kang and A. Kim wrote the manuscript.

